# Physical basis of lumen shape and stability in a simple epithelium

**DOI:** 10.1101/746792

**Authors:** Claudia G. Vasquez, Vipul T. Vachharajani, Carlos Garzon-Coral, Alexander R. Dunn

## Abstract

A continuous sheet of epithelial cells surrounding a hollow opening, or lumen, defines the basic topology of numerous organs. *De novo* lumen formation is a central feature of embryonic development whose dysregulation leads to congenital and acquired diseases of the kidney and other organs. Hydrostatic pressure has been proposed to drive lumen expansion, a view that is supported by recent experiments in the mouse blastocyst. High luminal pressure should produce lumen surfaces that bow outwards toward the surrounding cells. However, lumens formed in other embryonic tissues adopt highly irregular shapes, with cell apical faces that are bowed inward, suggesting that pressure may not be the dominant contributor to lumen growth in all cases. We used three-dimensional live-cell imaging to study the physical mechanism of lumen formation in Madin Darby Canine Kidney (MDCK) cell spheroids, a canonical cell-culture model for lumenogenesis. Our experiments revealed that neither lumen pressure nor the actomyosin cytoskeleton were required to maintain lumen shape or stability. Instead, we find that, in our model system, lumen shape results from simple geometrical factors tied to the establishment of apico-basal polarity. A quantitative physical model that incorporates cell geometry, cortical tension, and intraluminal pressure can account for our observations as well as cases in which pressure indeed plays a dominant role. Our results thus support a unifying physical mechanism for the formation of luminal openings in a variety of physiological contexts.

## Introduction

Lumens, or hollow openings surrounded by sheets of cells, are a ubiquitous structural feature of metazoans. While the molecular components required for lumen formation have been characterized in some detail, the physical mechanisms that underlie the initial steps in lumen formation remain less explored (Bryant et al., 2010; Cerruti et al., 2013; Ferrari et al., 2008; Martín-Belmonte et al., 2008; O’Brien et al., 2001; Odenwald et al., 2017; Rodríguez-Fraticelli et al., 2010; Sigurbjörnsdóttir et al., 2014; Wang et al., 1990). In this study we examined the mechanics of lumen formation and expansion in Madin Darby Canine Kidney (MDCK) cell spheroids, an archetypal cell culture model for studying lumenogenesis. Previous work in MDCK spheroids and other systems has assumed that lumen growth occurs due to intraluminal hydrostatic pressure (Chan et al., 2019; Dasgupta et al., 2018; Latorre et al., 2018; Ruiz-Herrero et al., 2017). In such a scenario, lumen shape and size are governed by the Young-Laplace equation, which states that the pressure difference (*P*) between the cells and the lumen is counterbalanced by the surface tension (*γ*) of the lumen surface (apical faces of the cells) and inversely proportional to the lumen radius (*r*): 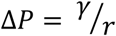. The presence of a luminal pressure is motivated by work that showed ion channels are critical for lumen formation and expansion *in vitro* and *in vivo* (Bagnat et al., 2007; Yap et al., 1993), and by pressure-driven fluctuations in lumen size in some model systems (Chan et al., 2019; Ruiz-Herrero et al., 2017).

Importantly, a positive luminal pressure should produce convex lumen surfaces that bow outwards toward the surrounding cells. In model systems such as the developing mouse blastocyst and the bile canaliculus this convex surface curvature is well-documented (Dasgupta et al., 2018; Dumortier et al., 2019). However, published images of some model systems, for example MDCK cell spheroids and various *in vivo* examples of lumens such as liver bile ducts, blood vessels, and pro-amniotic cavities, demonstrate areas of concave lumen curvature, where cell apical faces are bowed into the lumen (Benhamouche-Trouillet et al., 2018; Bryant et al., 2010; Cerruti et al., 2013; Ferrari et al., 2008; Hoijman et al., 2015; O’Brien et al., 2001; Oteiza et al., 2008; Shahbazi et al., 2016; Strilić et al., 2010; Wong et al., 2010). These observations suggest that a positive pressure gradient may not be the dominant contributor to the growth of all lumens. We therefore sought to understand the physical forces maintaining lumen shape in the context of *de novo* lumen formation.

## Results

### Lumens in small- and intermediate-size MDCK spheroids are irregularly shaped

We used MDCK cells as our model system to study lumen stability due to their ability to reliably establish apico-basal polarity and form lumens in 3D culture conditions in a manner that recapitulates lumenogenesis in *in vivo* model systems (Bryant et al., 2010; O’Brien et al., 2001; Wang et al., 1990). We seeded MDCK cells expressing a fluorescent marker for actin filaments (Lifeact-RFP) in the recombinant extracellular matrix Matrigel. Under these culture conditions, MDCK cells spontaneously form hollow spheroids within 24 hours. To obtain high-resolution images of nascent lumens, we imaged young (18-24 hours) spheroids using lattice light sheet microscopy (LLSM). This acquisition method produced images in which the two opposing apical cortices of the lumen are clearly distinguishable and separated by ∼200-300 nm (**Fig. 1a, b**). We infer that cellular apical surfaces are intrinsically non-adherent, as even small fluctuations in cell shape would allow apposing apical surfaces to contact and potentially adhere. This “non-stick” behavior may reflect the enrichment of negatively charged sialoglycoproteins such as Podocalyxin and/or the active exclusion of cell adhesion proteins (e.g. E-cadherin) (Ferrari et al., 2008; Kerjaschki et al., 1984; Strilić et al., 2010; Wang et al., 1990).

**Figure 1.**
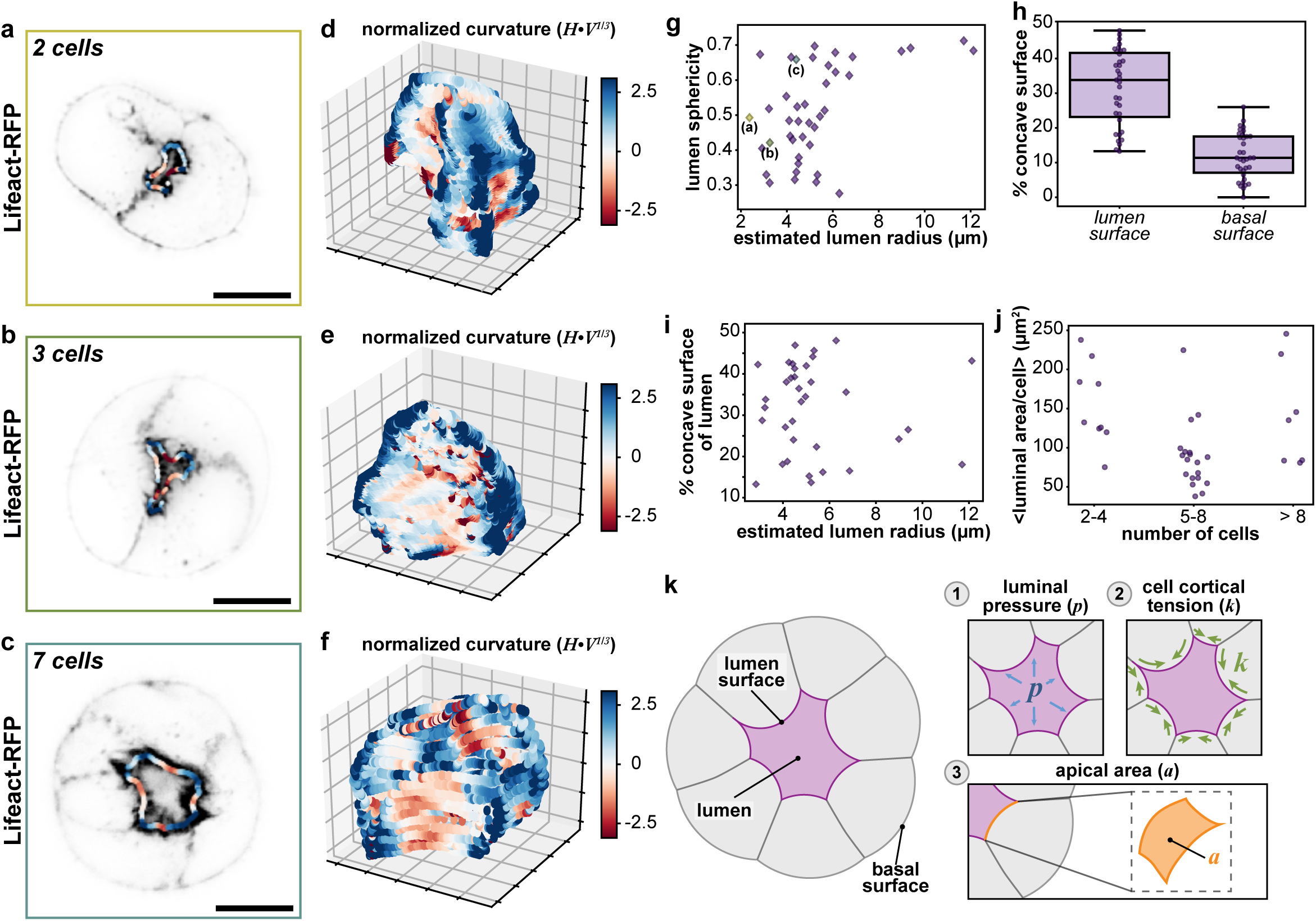
Quantification of lumen morphology. **a, b, c**, Representative single-plane images of MDCK spheroids expressing Lifeact-RFP (gray-scale). The mean lumen curvature is superimposed as a red-blue outline, where red is concave (negative local curvature) and blue is convex (positive local curvature). **d, e, f**, 3D contour plots of corresponding lumen surfaces showing local curvatures. **g**, Lumen sphericities plotted as a, function of estimated lumen radius. Estimated lumen radius (*r*) was calculated using lumen volume (*V*): *r = (3V/4π)*^*1/3*^. Values for representative spheroids from **a, b**, and **c** as indicated. **h**, Percent of lumen surface (left) and basal surface (right) that is concave (negative curvature). **i**, Percent of lumen surface that is concave (negative curvature) as a function of estimated lumen radius, as determined by lumen volume. **j**, Mean luminal surface area per cell plotted as a function of number of cells in the spheroid. **k**, Schematic depicting physical forces that could determine luminal shape. **(1)** luminal pressure (*p*) **(2)** cell cortical tension (*k*) and **(3)** apical area (*a*). Scale bars are 10 μm. Box plot in **h** shows median, quartiles of dataset, and whiskers extending to maximum and minimum of distributions. For plots **g-j, *n*** = 35 spheroids.

To derive insight into the physical mechanisms that dictate the shape and stability of small and intermediate-sized lumens, we used confocal microscopy to quantify the shapes of the luminal (apical) and outer (basal) surfaces of intermediate-size MDCK spheroids grown for 2 days in 3D conditions. As with two- to three-cell spheroids, the lumens of intermediate-sized spheroids (8-20 cells) were irregular in shape (**Fig. 1c, Supplementary Fig. 1a**). This observation is consistent with published shapes of MDCK lumens (Bryant et al., 2010; Cerruti et al., 2013; Ferrari et al., 2008; O’Brien et al., 2001) and other lumens observed in intact organisms (Benhamouche-Trouillet et al., 2018; Hoijman et al., 2015; Oteiza et al., 2008; Shahbazi et al., 2016; Strilić et al., 2010) though the irregularity of shape was not previously commented upon or explored in detail.

To determine how similar or dissimilar lumen shapes were to spheres, we calculated the sphericity *Ψ*, given by: 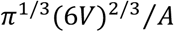, for lumen volume *V* and surface area *A*. This metric ranges from 0 (far from spherical) to 1 (exactly a sphere). The sphericity of lumens ranged from *Ψ*∼0.70 to very far from spherical (*Ψ*∼0.3) (**Fig. 1g**), values substantially less spherical than, for example, a cube (*Ψ* = 0.81). Further, we observed a lumen size-dependent crossover from irregular morphology to more spherical morphology: the lumens of large MDCK spheroids, with an estimated radius (calculated from lumen volume) of ∼10 µm and tens of cells, abruptly transitioned to more spherical shapes with *Ψ* ∼ 0.7.

In addition to lumen sphericity, we computed the mean curvature (***H***) at each voxel of lumen surface, normalized by lumen volume. These measurements demonstrate that lumens, even those with sphericity greater than 0.65, have areas of negative mean curvature, where the negative value denotes concave (inward) bending (**red; Fig. 1d-f, Supplementary Fig. 1b**). The percent of the total lumen surface area with negative mean curvature varied from 10% to over 50% (**Fig. 1h**). Unlike the trend for sphericity, there were no clear size-dependent trends towards a lower fraction of concavity as lumens grew larger (**Fig. 1i**). In contrast to the variability of lumen shape observed, the sphericity and percent concavity of the basal surfaces of the spheroids were close to 1 and 0%, respectively (**Fig. 1h, Supplementary Fig. 1c**).

These data are inconsistent with a model of positive luminal pressure as the sole driving mechanism to maintain the shape of small- and intermediate-sized lumens. Instead, we noted as well that the average luminal surface area per cell remained roughly constant for spheroids of 2 to >8 cells (**Fig. 1j**). This observation suggested an alternate scenario in which the apical surface area per cell was actively regulated, with apical surface area added faster than the equivalent amounts of luminal volume, thus leading to the observed irregular lumen shapes. Motivated by this possibility, we decided to more closely investigate three physical factors that could influence lumen shape: (1) intraluminal pressure, (2) cell cortical tension, and (3) a preferred apical domain size (**Fig. 1k**).

### Intraluminal pressure does not significantly define lumen shape or stability

To test how modulating intraluminal pressure affected lumen shape, we treated MDCK spheroids grown in Matrigel for either 3 or 7 days with small molecules that act to promote or inhibit apical fluid secretion: the V2 receptor agonist 1-desamino-8-D-AVP (ddAVP, 10µM) and the Na^+^/K^+^ ATPase inhibitor ouabain (330 µM), respectively (Chan et al., 2019; Grant et al., 1991; Schoner, 2002) (**Fig. 2a**). Treatment with ddAVP for 4 hours caused lumen cross-sectional area to increase compared to vehicle controls while having little change in cell thickness, consistent with the expected increase in lumen volume (**Fig. 2b, c, Supplementary Fig. 2a, b**). We also observed an unanticipated, modest increase in lumen cross-sectional area for spheroids treated with ouabain, though only for 7-day old (larger) spheroids (**Supplementary Fig. 2c**).

**Figure 2.**
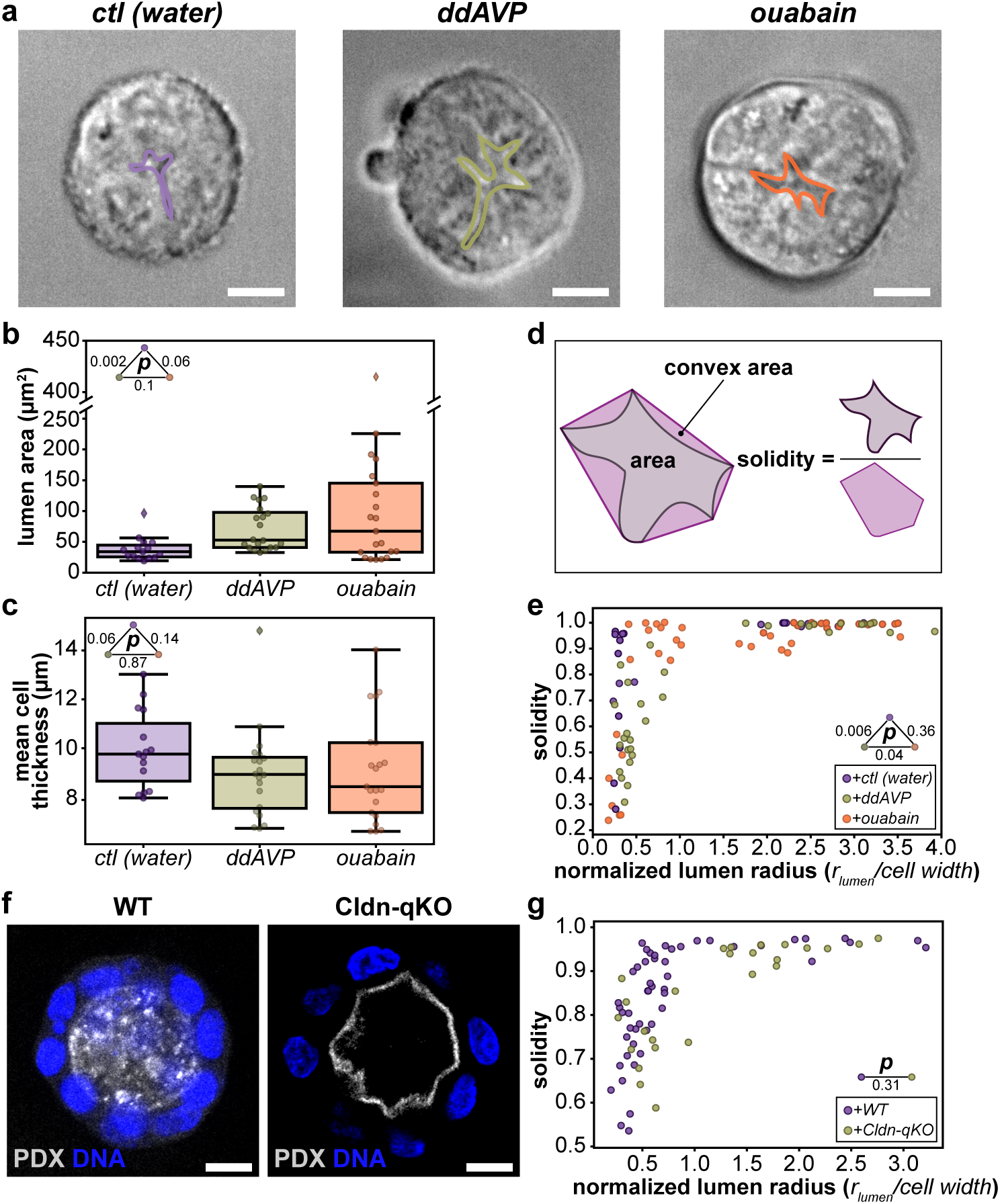
Manipulation of intraluminal pressure does not significantly affect lumen shape. **a**, Representative cross-sections of MDCK spheroids grown for 3 days imaged using differential interference contrast treated with vehicle (left), ddAVP (middle), or ouabain (right) for 4 hours. Lumens are outlined in purple, green, and orange, respectively. **b**, Quantification of cross-sectional luminal area of MDCK spheroids grown for 3 days in control, ddAVP, and ouabain conditions (p-values are from rank-sum test). **c**, Quantification of mean cross-sectional cell thickness of MDCK spheroids grown for 3 days in control, ddAVP, and ouabain conditions (p-values are from rank-sum test). **d**, Schematic of calculating solidity, a metric that reflects surface irregularity. **e**, Solidity of lumens from MDCK spheroids grown for 3 days and 7 days, treated as indicated with vehicle (purple), ddAVP (green), or ouabain (orange) for 4 hours as a function normalized lumen radius (ratio of estimated lumen radius determined by lumen volume and mean cell width) (p-values from 2D Kolmogorov-Smirnov test). **f**, Representative images of fixed wildtype (WT, left) and Claudin-quintuple KO (Cldn-qKO, right) MDCK spheroids with lumens immunostained for the apical surface protein podocalyxin (PDX, white) and DNA (blue) (Otani et al., 2019). **g**, Lumen solidity plotted as a function of normalized lumen radius for wildtype lumens (WT, purple), Cldn-qKO lumens (green) (p-value from 2D Kolmogorov-Smirnov test). Scale bars are 10 μm. Box plots in **b** and **c** show median, quartiles of dataset, and whiskers extending to maximum and minimum of distributions, excluding outliers (indicated with diamonds). For plots **b** and **c, *n*** = 15, 21, 20 spheroids for control, ddAVP, and ouabain conditions, respectively. For plot **e, *n*** = 48, 33, 42 spheroids for control, ddAVP, and ouabain conditions, respectively. For plot **g, *n*** = 51 and 25 spheroids for WT and Cldn-qKO conditions, respectively.

To determine how these treatments altered lumen shape, we calculated the solidity of lumen cross-sectional shapes. The solidity metric is defined as the ratio of the lumen area and the convex area (i.e., convex hull of the lumen shape) (**Fig. 2d**). A value of 1 describes completely convex shapes, such as an ellipse or a regular polygon, while lower values reflect varying degrees of surface irregularity. Independent of treatment with ddAVP, ouabain, or water vehicle control, the solidity of lumens showed the same abrupt transition to values close to 1 with increasing lumen size (**Fig. 2e**). Although control and ddAVP treated lumens were statistically distinguishable by a 2D Kolmogorov-Smirnov test, differences in lumen shape were, in practice, not readily discerned (**Fig. 2a, e**). Thus, for lumens of approximately four to tens of cells, modulating pressure did not strongly influence lumen shape.

To further probe this observation, we measured the solidity of lumens formed by spheroids grown from either wildtype MDCK cells or an MDCK cell line lacking the five most highly expressed claudins (quintuple knockout; Cldn-qKO) (Otani et al., 2019). Claudins are essential components of the tight junction. A previous study showed that monolayers formed by Cldn-qKO cells showed dramatically increased small-molecule permeability relative to wildtype cells. It is therefore unlikely that Cldn-qKO cells could maintain a pressure differential between the lumen and the exterior environment. Remarkably, but consistent with prior reports, Cldn-qKO cells formed lumens (Otani et al., 2019) (**Fig. 2f**). Moreover, lumens formed by wildtype and Cldn-qKO cells followed the same relation between normalized lumen radius and solidity (**Fig. 2g**). We conclude that, at least in this model system, pressure alone is not a driver for determining lumen shape and stability.

### Cell cortical tension subtly influences lumen volume but not shape or stability

Cortical tension helps define the shape of cells and could thus be an important modulator of lumen shape and stability. We experimentally decreased cell cortical tension with a cocktail of inhibitors (1 µM latrunculin A, 20 µM ML-7, 20 µM Y-27632, and 50 µM nocodazole) that acutely ablates the actomyosin and microtubule cytoskeletons (Owen et al., 2017) (**Fig. 3a, b, Supplementary Video 1**). This test also served to determine if intraluminal pressure significantly stabilized lumen shape: if the lumen were under positive pressure, softening the cell cortices would cause the lumen to become rounder, as the luminal pressure would push the apical surfaces outwards. On the whole, treatment resulted in small increases, on average, in luminal volume (**Fig. 3c**), though without a significant change in lumen surface area (**Fig. 3d**). Notably, this treatment did not significantly affect the volume of the whole spheroid, indicating that in this observed time frame the environment surrounding the spheroid does not collapse in (**Supplementary Fig. 3**). Treatment likewise did not obviously influence lumen solidity (**Fig. 3e**). Thus, cytoskeletal ablation led to only subtle changes in lumen size and shape and did not alter lumen stability.

**Figure 3.**
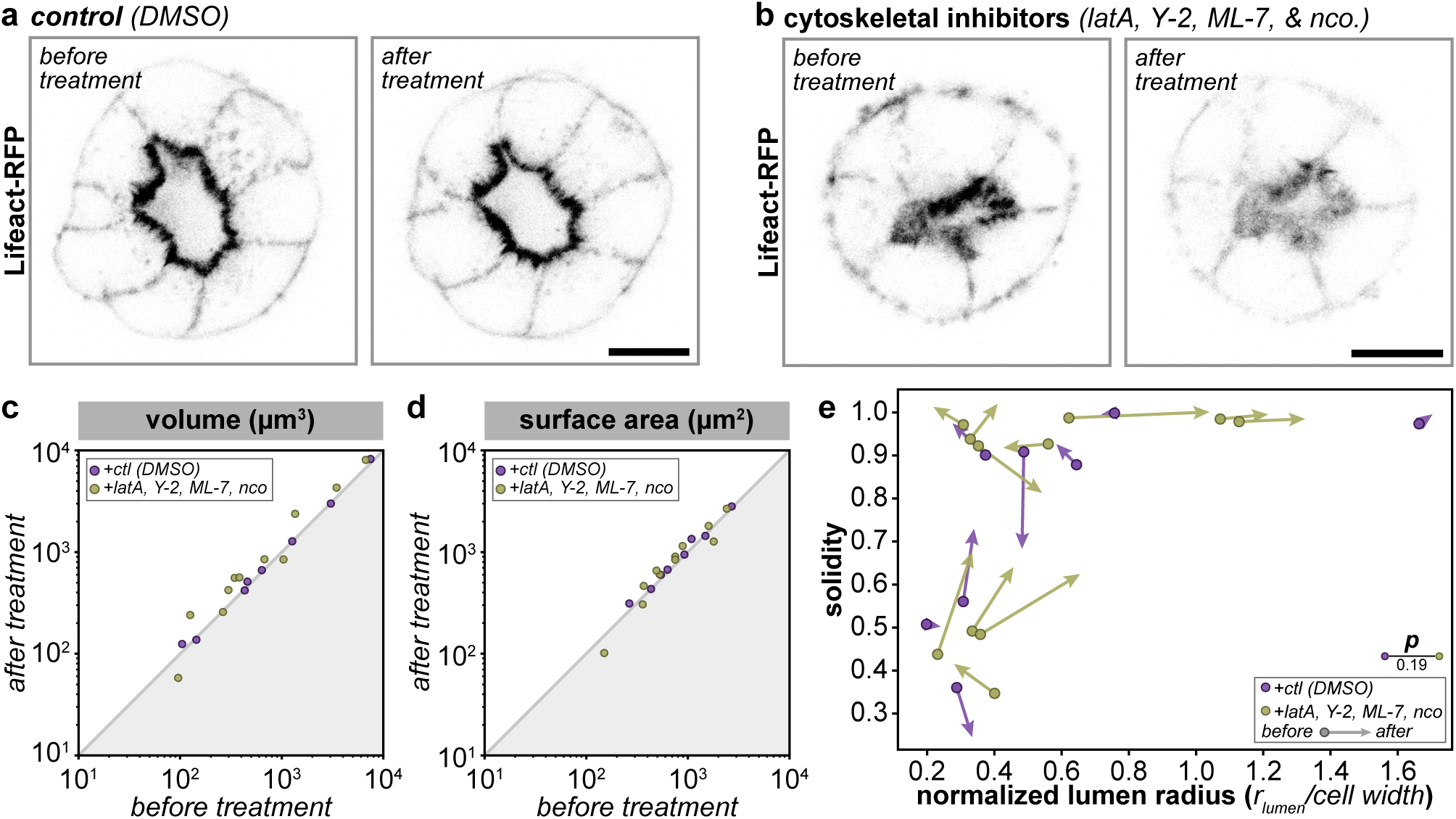
Acute cytoskeletal ablation does not significantly alter lumen shape. **a**, Representative single-plane images MDCK spheroids expressing Lifeact-RFP 2 min before (left) and 18 min after (after) addition of DMSO vehicle control. **b**, Representative single-plane images MDCK spheroids expressing Lifeact-RFP 2 min before (left) and 18 min after (right) treatment with a cytoskeletal inhibitor cocktail (latrunculin A, Y-27632, ML-7, and nocodazole). **c**, Log-log plot of lumen volume before (*t* = −2 min) and after treatment (*t* = +18 min) (rank-sum test DMSO-cytoskeletal inhibitors p = 0.13, Wilcoxon signed rank test *Δt*(DMSO) p = 0.263, *Δt*(cytoskeletal inhibitors) p = 0.041). **d**, Log-log plot of lumen surface area before (*t* = −2 min) and after treatment (*t* = +18 min) (rank-sum test DMSO-cytoskeletal inhibitors p = 0.41, Wilcoxon signed rank test *Δt*(DMSO) p = 0.069, *Δt*(cytoskeletal inhibitors) p = 0.091). **e**, Lumen solidity plotted before (arrow end) and after (arrowhead) treatment with DMSO vehicle control (purple) or cytoskeletal inhibitor cocktail (green) as a function of normalized lumen radius (p-values from 2D Kolmogorov-Smirnov test). Scale bars are 10 µm. For plots **c-e, *n*** = 8 and 11 spheroids for DMSO and cytoskeletal inhibitors conditions, respectively.

### Modulation of apical area alters lumen shape

To test our intuition that the size of the apical domain influences lumen shape we searched the literature for manipulations that drive the expansion of the apical domain. Apical membrane identity is determined by two complexes: the Crumbs complex and the Par complex (Bryant and Mostov, 2008; Riga et al., 2020). Crumbs3a, a Crumbs homolog, is critical for establishment of apical domains and consequently lumen formation in MDCK spheroids (Schlüter et al., 2009). Overexpression of Crumbs3a resulted in apical membrane expansion into the lumen, even when lumens were quite large (estimated radius ∼ 20 µm) (Schlüter et al., 2009). The Par complex is composed of the polarity proteins Par3 and Par6 and the atypical kinase aPKC. Tight regulation of aPKC activity is necessary for proper polarity establishment and maintenance. The membrane-associated protein, KIBRA, an upstream regulator of the YAP/TAZ pathway, can bind to aPKC and inhibit its activity (Yoshihama et al., 2011). Consequently, knockdown of KIBRA also results in expansion of the apical domain into the lumen (Yoshihama et al., 2011). We measured the shapes of published examples of lumens produced by MDCK cells overexpressing Crumbs3a (Schlüter et al., 2009) or with KIBRA knockdown (Yoshihama et al., 2011). In published examples, these manipulations led to a marked decrease in lumen solidity, even at large lumen sizes (**Fig. 4a**). Thus, apical domain size could markedly influence lumen shape.

**Figure 4.**
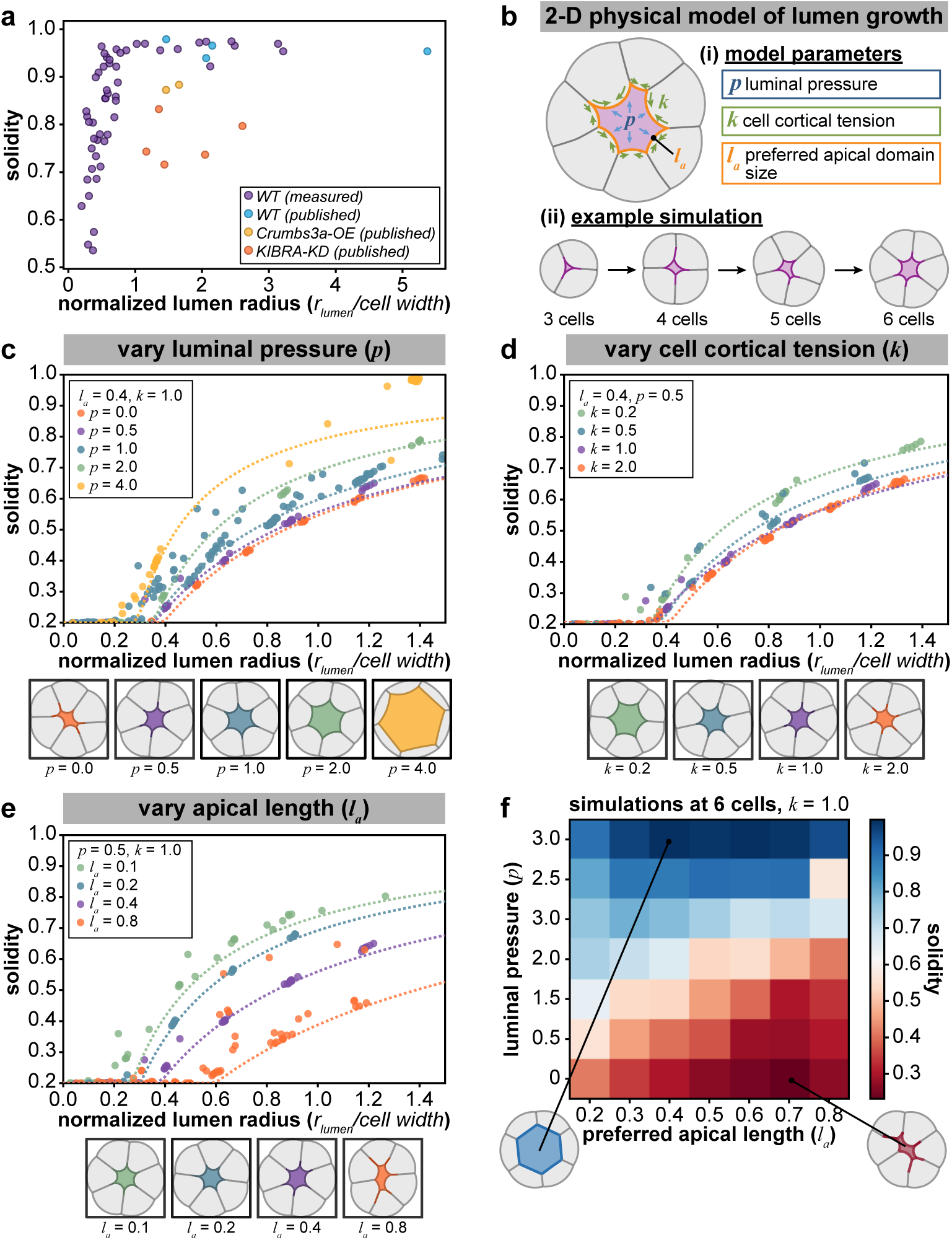
Size of apical domain controls lumen shape more acutely than luminal pressure or cortical cell contractility. **a**, Lumen solidity plotted as a function of normalized lumen radius for wildtype (WT, measured, purple and published, blue) and published apical expansion manipulations (Crumbs3a overexpression (Crumbs3a-OE), yellow and KIBRA knockdown (KIBRA-KD), orange). **b**, Schematic depicting parameters of vertex-like model describing lumen shape **(i)**, and example simulation showing spheroid growth from 3 to 6 cells **(ii). c-e**, Plots of solidity as a function of normalized lumen radius from simulations of 2D vertex-like model (top) and representative outputs of model grown to 6 simulated cells (bottom), keeping preferred apical length and cell cortical tension fixed (*l*_*a*_ = 0.4, *k* = 1.0) while varying luminal pressure (*p* = 0.0-4.0) **(c)**, keeping preferred apical length and luminal pressure fixed (*l*_*a*_ = 0.4, *p* = 0.5) while varying cell cortical tension (*k* = 0.0-2.0) **(d)** and keeping luminal pressure and cell cortical tension fixed (*p* = 0.5, *k* = 1.0) while varying preferred apical length (*l*_*a*_ = 0.1-0.8) **(e). f**, Heatmap of mean solidity for outputs of simulations while varying luminal pressure and preferred apical length. Representative outputs of maximum (blue) and minimum (red) solidity reached in simulations. For plot **a, *n*** = 51, 4, 2, 5 spheroids for WT, WT (published), Crumbs3a-OE (Schlüter et al., 2009), and KIBRA-KD (Yoshihama et al., 2011) conditions, respectively.

### A minimal model of preferred apical domain size can explain features of lumen geometry

The data above suggested that apical domain size, rather than pressure or cortical tension, might play a dominant role determining lumen shape. We sought to develop a physical model that could account for our observations. Although such mathematical models cannot be proven to be correct, they are a useful means of testing the underlying biological model against available data, and of making predictions to guide future experiments.

We adapted a vertex-based model of tissue mechanics to construct a simple two-dimensional model of a growing spheroid (Fletcher et al., 2017; Polyakov et al., 2014). This model incorporates only intraluminal pressure (***p***), cell cortical tension (***k***), and preferred apical domain size (***l***_***a***_) (**Fig. 4b**). Spheroids were simulated as they grew from 3 to 10 cells in size for each set of parameters. Within this model, we systematically varied ***p, k***, and ***l***_***a***_, and quantified lumen shape using solidity as a metric. As expected, increasing ***p*** or decreasing ***k*** led to modest increases in solidity at a given relative lumen size (**Fig. 4c, d**). Importantly, lumens tended to have higher solidity with increasing size even if luminal pressure was zero (**Fig. 4c, teal**), an outcome in line with experimental observation (**Fig. 2g**). Further, increasing ***p*** or decreasing ***k*** modestly affected lumen size and solidity when the simulations had fewer than 5 cells; however, as the number of cells increased, a large positive pressure or small cortical tension resulted in larger and more regular-shaped lumens (**Supplementary Fig. 4a-d, green**). These trends agree with the modest increases in lumen volume observed upon increased lumen pressure (**Fig. 2b**) or cytoskeletal ablation (**Fig. 3c**). Increasing ***l***_***a***_ (preferred apical domain size) led to marked decreases in solidity (**Fig. 4e**), in agreement with our quantification of literature data (**Fig. 4a**). Thus, lumen shape was more sensitive to alterations in apical domain size than to comparable fold-changes in cortical tension or intraluminal pressure, in line with our experimental results. An important prediction of the model is that preferred apical domain size and pressure together set the lumen size at which this crossover to round lumens occurs. Thus, lumen shape can be stabilized and regularized by pressure-dependent and -independent mechanisms depending on the biological system (**Fig. 4f**).

While the model incorporates pressure, tension, and preferred apical domain size parsimoniously, it is incomplete in several ways. First, it models a two-dimensional cross-section of a three-dimensional object. Second, it does not account for differences in cortical tension between apical and basolateral domains, or differences in physical properties between different cells within the same spheroid. Finally, it does not provide details as to the mechanism by which apical domain size might be regulated. Nonetheless, its predictions qualitatively agree with our data in ways that pressure-only models cannot. In particular, this minimal model is sufficient to capture the most notable trends in our data, and others, regarding lumen shape and size (Chan et al., 2019; Dumortier et al., 2019; Latorre et al., 2018; Ruiz-Herrero et al., 2017).

## Discussion

Our findings highlight a pressure-independent method for stabilizing lumens that, to our knowledge, has been largely overlooked despite its probable prevalence (Alvers et al., 2014; Dong et al., 2014; Gao et al., 2017; Shahbazi et al., 2016; Tsarouhas et al., 2007). Current evidence demonstrates that hydrostatic pressure plays a central role in the growth and stabilization of lumens in some circumstances, most notably the mammalian blastocyst. Our data indicate that lumens can also expand via a distinct, pressure-independent pathway in which lumen growth occurs by the addition of a roughly constant amount of apical membrane per cell, with sufficient fluid transport to allow the lumen to gradually increase in volume while avoiding large positive or negative pressures. The advantages of pressure-independent lumen growth remain to be firmly established. However, we note that this mechanism is predicted to be physically robust and energy efficient relative to pressure-driven growth, which exposes tissues to pressure-driven rupture (Chan et al., 2019; Ruiz-Herrero et al., 2017).

In order to better understand lumen growth, we developed a physical model that can account for our data as well as pressure-driven and -independent lumenogenesis in a wide variety of model systems. In this model, the creation of a non-stick apical membrane, which occurs via directed vesicular trafficking (Bryant and Mostov, 2008; Chan and Hiiragi, 2020; Sigurbjörnsdóttir et al., 2014), is itself sufficient to define the contours of the lumen. Lumen shape is however dictated by the balance of preferred apical area, pressure, and cortical tension, with high pressure, high cortical tension, and small apical domains yielding regular shapes. This model can account for the observation of the large variety of both regular and irregular lumen shapes that have been described in different model systems. Pressure-dependent and pressure-independent mechanisms for dictating lumen shape thus coexist and fulfill complementary functions in driving embryonic growth and tissue morphogenesis.

Previous work demonstrates that the establishment of apico-basal polarity and the earliest stages in lumen formation are tightly coupled at a molecular level (Bryant et al., 2010; Bryant and Mostov, 2008; O’Brien et al., 2001; Sigurbjörnsdóttir et al., 2014). Our work builds on these observations and highlights the deep connections between the establishment of a distinct, lumen-facing apical membrane and the physical mechanisms that stabilize small lumens. The creation of hollow lumens is likely to be an evolutionarily ancient innovation that was key to the construction of multicellular tissues (Anderson et al., 2017; Brunet et al., 2019; Yin et al., 2018; Yin et al., 2015). Evolutionary data indicate the molecular components required for the establishment of a defined apical domain are likewise ancient, and appeared simultaneously with the advent of multicellular animals (Belahbib et al., 2018). We speculate that the evolutionary origins of apico-basal polarization and of lumenogenesis may be inextricably linked, and that their joint appearance constituted a key evolutionary innovation enabling the construction of animal life.

## Acknowledgements

C.G.V. is supported by the NIH NIGMS (1F32GM125113-01). V.T.V. is supported by the Stanford Medical Scientist Training Program (NIH T32GM007365). C.G.C is supported by a long-term postdoctoral fellowship from the Human Frontier Science Program. A.R.D. acknowledges the HHMI (Faculty Scholar Award), and the NIH (R01GM117457, R35GM130332). We want to thank Dr. Christina Hueschen, other members of the Dunn lab, and Dr. Lucy E. O’Brien for discussion and insightful comments on the manuscript. We want to thank Dr. Tetsuhisa Otani and Dr. Mikio Furuse for their generous gift of the MDCK Claudin-quintuple knockout cell line.

## Author contributions

C.G.V., V.T.V, and C.G.C. performed experiments and analyzed data. V.T.V developed the models with input from C.G.V. and A.R.D. All authors wrote and edited the manuscript.

## Methods

### Cells culture and generation of cell lines

MDCK II (Sigma Cat. #00062107) cells were cultured at 37 °C and 5% CO_2_ in DMEM (Thermo Fisher Cat. #11885076) supplemented with 10% fetal bovine serum (FBS, Corning) and 1% penicillin-streptomycin (ThermoFisher). Live-cell confocal and brightfield microscopy experiments were performed in Leibovitz’s L15 media (L15, ThermoFisher) supplemented with 10% FBS (Corning) and 1% penicillin-streptomycin. Live-cell lattice light sheet microscopy was performed in L15 media supplemented with 1% FBS and Insulin-Transferrin-Selenium (Invitrogen). MDCK cells constitutively expressing Lifeact-RFP were generated to visualize the actin cytoskeleton. Briefly, cells were transfected with a plasmid containing the PiggyBac transposon system and the Lifeact-RFP sequence (DNA2.0), cells were selected for plasmid integration with geneticin (G418, ThermoFisher). MDCK Claudin-quintuple knockout (Cldn-qKO) cells were a generous gift from Tetsuhisa Otani and Mikio Furuse (National Institute for Physiological Science, Japan) (Otani et al., 2019).

### Generating MDCK spheroids

MDCK spheroids were generated as previously described in (Peterman and Prekeris, 2017; Vieira et al., 2006). Briefly, 75 µL of cell suspension containing ∼10^4^ cells were mixed with 150 µL Matrigel GFR (Corning CB-40230). 25 µL drops of cell-Matrigel suspensions were seeded into each well of an 8-well chambered coverglass (Nunc, No. 1.5) and incubated at 37 °C for 30 minutes before adding DMEM to cells. Media was changed every other day. At least 2 hours prior to live-imaging experiments, media was changed to L15 with supplements (as indicated above).

### Pharmacological inhibition

To disrupt ion and fluid pumping, spheroids were treated with ouabain, a Na^+^/K^+^ ATPase inhibitor (Sigma) or 1-deamino-8-D-arginine vasopressin (ddAVP, Sigma), a vasopressin receptor agonist. Each was dissolved in distilled water at 1000X final concentration and diluted in cell culture media immediately before treatment for 4-24 hours before DIC imaging. Ouabain was used at a final concentration of 333 µM while ddAVP was used at a final concentration of 10 µM. For cytoskeletal inhibition (and controls) experiments, data were collected on at least 5 separate days from distinct samples. For fluid pumping inhibition experiments, data were collected on two separate days from distinct samples. To perturb the actomyosin and microtubule cytoskeletons, cells were treated with a cytoskeletal inhibitor cocktail composed of latrunculin A (Sigma, 1 µM), Rho-kinase inhibitor Y-27632 (STEMCELL Technologies, 20 µM), MLCK inhibitor ML-7 (Enzo, 20 µM), and nocodazole (Sigma, 50µM). The cytoskeletal inhibitor cocktail was made at 5X final concentration in L15 media and 100 µL were added to imaging well with 400 µL of L15.

### Cell immunofluorescence

MDCK cells were fixed with 4% paraformaldehyde in PBS for 15 min at room temperature. Samples were blocked and permeabilized in 0.1% Triton, 1% bovine serum albumin (BSA, Sigma) in PBS for 1 h at room temperature. Cells were incubated in primary antibody solution in 0.1% Triton, 1% BSA in PBS overnight at 4 °C, and in secondary antibody solution in 0.1% Triton, 1% BSA in PBS for 2 hours at room temperature. To identify nuclei, Hoechst solution (Hoechst 34580, Invitrogen) was added at 1:1000 dilution to secondary antibody solution. Antibodies and corresponding concentrations used in this investigation are listed in **Table S1**. Images were acquired at room temperature (∼22 °C) using an inverted Zeiss LSM 780 confocal microscope with a 40X/1.3 NA C-Apo water objective, 405 nm diode laser, 561 nm diode laser, 633 nm HeNe laser, and a pinhole setting between 1 and 2 Airy Units. All images were acquired using Zen Black software (Carl Zeiss).

**Table S1.** Antibodies used and concentrations.

| Antibody | Dilution | Source |
| --- | --- | --- |
| mouse anti-gp135 (PDX) | 1:50 | Developmental Studies Hybridoma Bank (3F2/D8) |
| goat anti-mouse IgG Alexa Fluor 555 | 1:1000 | Cell Signaling Technology (Cat. 4413) |
| goat anti-mouse IgG Alexa Fluor 647 | 1:1000 | Cell Signaling Technology (Cat. 4410) |

### Lattice light sheet microscopy

We used a custom-built lattice light sheet microscope (LLSM) (Chen et al., 2014) to image MDCK spheroids. Spheroids were grown in 3 μL droplets of Matrigel without Phenol red (Corning) seeded on top of a 5 mm round cover glass (Warner Instruments). The samples were incubated for 12-36 hours at 37 °C in 25 mm tissue culture plates. Before experiments, samples were transferred to LLSM imaging medium (L15 media supplemented with 1% FBS and Insulin-Transferrin-Selenium Invitrogen) for 12-16 hours. Data were collected on two separate days from distinct samples.

Samples were illuminated by a 561 nm diode laser (0.5W, Coherent) using an excitation objective (Special Optics, 0.65 NA with a working distance of 3.74 mm) at 2% AOTF transmittance and laser power of 100 mW. Order transfer functions were obtained empirically by acquiring point-spread functions using 200 nm TetraSpeck beads adhered freshly to 5 mm glass coverslips (Invitrogen T7280) for each wavelength and for each day of experiments.

To achieve structured illumination, a square lattice was displayed on a spatial light modulator. This lattice was generated by an interference pattern of 59 Bessel beams separated by 1.67 µm and cropped to 0.22 with a 0.325 inner NA and 0.40 outer NA. The lattice light sheet was dithered 25 µm to obtain homogeneous illumination with 5% flyback time. Fluorescent signal was collected by a Nikon detection objective (CFI Apo LWD 25XW, 1.1 NA, 2 mm working distance), coupled with a 500 mm focal length tube lens (Thorlabs), a Semrock filter (BL02-561R-25) and sCMOS cameras (Hamamatsu Orca Flash 4.0 v2) with a 103 nm/pixel magnification.

Z-Stacks were acquired by moving the lattice light sheet and the detection objective synchronously, using a galvo mirror coupled at the back focal plane of the illumination objective and a piezomotor, respectively. The slices of the stacks were taken with an interval of 100 nm through ranges of 30-35 μm at 100ms camera exposure with 1-5 second intervals between z-stacks.

Raw data was flash corrected (Liu et al., 2017) and deconvolved using an iterative Richardson-Lucy algorithm (Chen et al., 2014) run on two graphics processing units (NVIDIA, GeForce GTX TITAN 4Gb RAM). Flash calibration, flash correction, channel registration, order transfer function calculation and image deconvolution were done using the LLSpy open software (Lambert and Shao, 2019). Visualization of the images and volume inspection were done using Spimagine (https://github.com/maweigert/spimagine) and ClearVolume (Royer et al., 2015).

### Confocal microscopy

All live confocal images were acquired at 37 °C using an inverted Zeiss LSM 780 confocal microscope with a 40X/1.3 NA C-Apo water objective, 561 nm diode laser, and a pinhole setting between 1 and 2 Airy Units. All images were acquired using Zen Black software (Carl Zeiss).

### Differential interference contrast (DIC) microscopy

DIC microscopy was performed at 37 °C using a Nikon Ti-E inverted microscope with a 20X/0.5 NA Plan Fluor CFI objective and N2 DIC prisms. Image acquisition was controlled using micromanager software (Edelstein et al., 2014).

### Image analysis

A Fiji (Schindelin et al., 2012) plugin was used to correct 3D drift in all live-cell confocal images (Parslow et al., 2014; Schindelin et al., 2012).

#### Cell number

The number of cells in each spheroid were manually and independently determined by CGV and VTV using the Lifeact-RFP signal to determine cell-cell boundaries.

#### Segmentation and surface shape calculations

Custom-developed Python code was used to detect and segment the lumens (i.e. apical surface) and basal surfaces spheroids. Briefly, lumen and basal surface boundaries were detected in each slice from segmented images using OpenCV. The boundary was parameterized by contour length. *X*- and *Y*-coordinates of each boundary were fit to Fourier series of varying order up to 15 (Schmittbuhl et al., 2003). A Bayesian Information Criterion was used to select the Fourier order to minimize overfitting (Neath and Cavanaugh, 2012). The smoothed boundary was used for calculations of local curvature, volume, and surface area. From these calculations sphericity (/) was computed as:

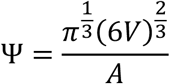

Where ***V*** is surface volume, and ***A*** is surface area.

Estimated lumen radius was calculated from the lumen volume as:

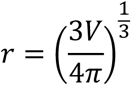

Where ***V*** is lumen volume.

#### Mean curvature

For each voxel on a surface, the smoothed contours in *XY* and *YZ* correspond to 1-dimensional curves on the surface, which are orthogonal at that voxel. The curvatures of these curves were computed using the Fourier representation. From these two curves, the surface mean curvature was estimated as follows: the surface normal vector was estimated as the cross product of the unit tangent vectors to each of these orthogonal cross-sectional curves. Because these two cross-sections are orthogonal, this is an appropriate approximation. Let a cross-section contour be given by 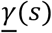. The Frenet-Serret formula gives 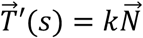, where 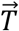 and 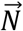 are the unit normal and binormal vectors of the contour, respectively. The curvature can be related to the geodesic (*k*_*g*_) and normal (*k*_*n*_) curvatures by the formula 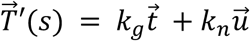, where 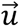 is the surface unit normal vectors, and 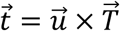, which also follows from the Frenet-Serret formula (Gray, 1997). The normal curvatures of each cross-sectional curve were thus computed from the estimated surface normal vector and the Frenet-Serret normal vector of that curve as 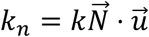. The mean curvature was then calculated as the mean of the two orthogonal normal curvatures.

#### Determination of fraction concave surface

Each segmented surface (lumen or basal surface) has a distribution of mean curvatures. To calculate the fraction concave surface area, we determined what fraction of each surface had negative (concave) mean curvatures.

#### Determination of mean luminal area per cell

Each segmented lumen surface was divided by the number of cells in the spheroid.

#### DIC image analysis

For each spheroid imaged in DIC, the apical (lumen) and basal surfaces were traced manually in Fiji (Schindelin et al., 2012) using the polygon selection tool. For apical (***a***)and basal (***b***) surfaces, the enclosed area (***A***) and perimeter (***P***) were measured automatically in Fiji. Mean cell thickness was computed as:

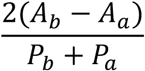

#### Determination of solidity

Solidity metric was calculated using Fiji plugin to measure the calculate and determine the area of the convex hull (***A***_***c***_) of the lumen shape. Solidity was computed by:

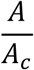

A solidity of 1 describes a completely convex shape, such as an ellipse or a regular polygon, while lower values reflect varying degrees of su<u>rfa</u>ce irregularity. For 3D confocal datasets, middle XY-plane of lumen was manually chosen for solidity analysis.

#### Determination of normalized lumen radius

For each spheroid, the lumen and spheroid radius (***r***) were determined calculated from the lumen area a<u>n</u>d spheroid area (***A***), respectively, as follows:

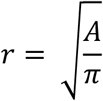

The normalized lumen radius (***r***_***norm***._) was calculated from these estimated radii as follows:

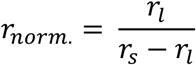

Where ***r***_***l***_ is the estimated lumen radius and ***r***_***s***_ is the estimated spheroid radius.

### Vertex-based model

We adapted a vertex-based model of tissue mechanics to construct a simple two-dimensional model. This model incorporates only a preferred apical area, cell cortical tension, and the geometric constraints of a spheroid.

Each cell in an *N*-cell model spheroid was described by four boundaries: a curved basal boundary, two straight lateral boundaries, and a curved apical boundary with preferred length *l*_*a*_. These boundaries have lengths and enclose a cell area *A*. Thus, the spheroid shape is completely determined by the locations of the apical and basal vertices, and the curvatures of the apical and basal boundaries. An *N*-cell spheroid was thus represented as a *6;*-dimensional vector which records the *x,y* positions of each vertex, the apical curvature, and the basal curvature.

For simplicity, we assume that the average area of a cell in the spheroid is constant. The effects of cortical tension at the basal and lateral boundaries, and preferred apical domain size, are modeled as springs with rest lengths of 0, 0, and *l*_*a*_, respectively. Finally, we include an energetic term that favors higher luminal areas, parametrized by a pressure difference between cells and the lumen *P*_*L*_, which may be set to zero.

Thus, in line with previous descriptions of such vertex-based models (Polyakov et al., 2014; Trushko et al., 2020) the energy of the system is given by:

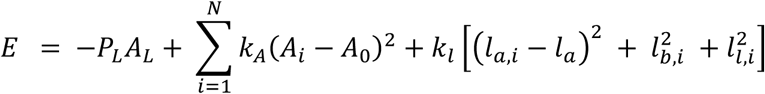

This can be nondimensionalized by dividing by the characteristic energy 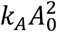, which yields the dimensionless equation:

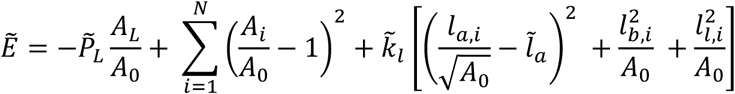

where we have the following three dimensionless parameters:

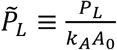, a dimensionless pressure

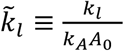, a dimensionless cortical tension

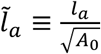, a dimensionless preferred apical domain size.

This nondimensionalized equation was used to simulate a growing spheroid as follows:

1. A 3-cell spheroid was generated with basal vertices evenly-spaced around a circle of radius 2; apical vertices chosen independently, uniformly at random from the interior of the unit circle; and apical curvatures chosen uniformly at random from the interval [1,2].
2. Stochastic gradient descent was used to minimize the energy of this spheroid such that all vertices were within 0.01 dimensionless units of a local minimum.
3. The highest-energy cell was divided by adding an additional apical and basal vertex in the center of the apical and basal boundaries, respectively.
4. The resulting 4-cell spheroid was again energy-minimized using stochastic gradient descent.
5. The process was repeated up to 10-cell spheroids.

Simulations were performed using Python using the NumPy and SciPy libraries.

### Statistical analysis

Statistically significant differences between control and drug treatment groups were assessed via a Rank-sum test, as indicated (**Fig. 2b, c, 3c,d, Supplementary Fig. 2 and 3**). Statistically significant differences between trends of control and drug or wildtype and Cldn-qKO groups were assessed via two-dimensional Kolmogorov-Smirnov test. Data comparing before and after treatment were assessed using the Paired Wilcoxon test (**Fig. 3c, d, Supplementary Fig. 3**). Rank-sum and Paired Wilcoxon test statistical analyses were performed using the stats module of the SciPy Python package. Two-dimensional Kolmogorov-Smirnov tests was implemented in Python as described in (Peacock, 1983).

### Code availability

Python analysis procedures are available from the corresponding author upon request.

### Data availability

All data generated or analyzed in this study are available from the corresponding author upon request.

---

**Supplementary Figure 1.**
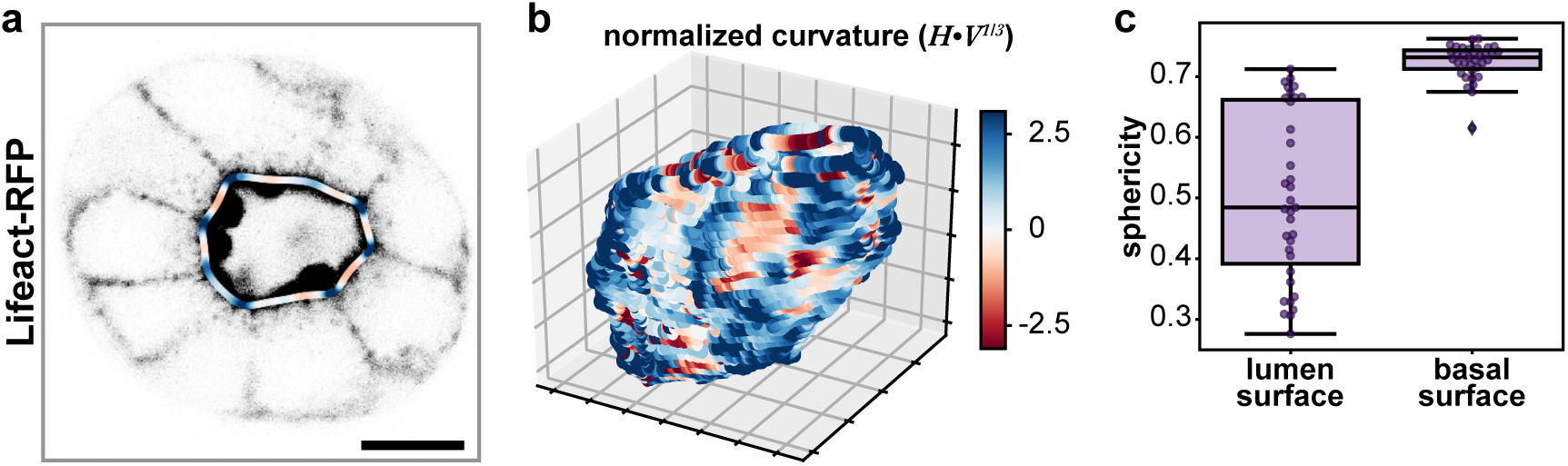
**a**, Representative single-plane of an MDCK spheroid expressing Lifeact-RFP (gray). The mean lumen curvature is superimposed as a red-blue outline, where red is concave (negative local curvature) and blue is convex (positive local curvature). **b**, 3D contour plots of corresponding lumen surfaces showing local curvatures. **c**, Sphericity lumen surface (left) and basal surface (right) (***n*** = 35 spheroids). Scale bars are 10 μm. Box plot in **c** shows median, quartiles of dataset, and whiskers extending to maximum and minimum of distributions, excluding outliers (indicated with diamonds).

**Supplementary Figure 2.**
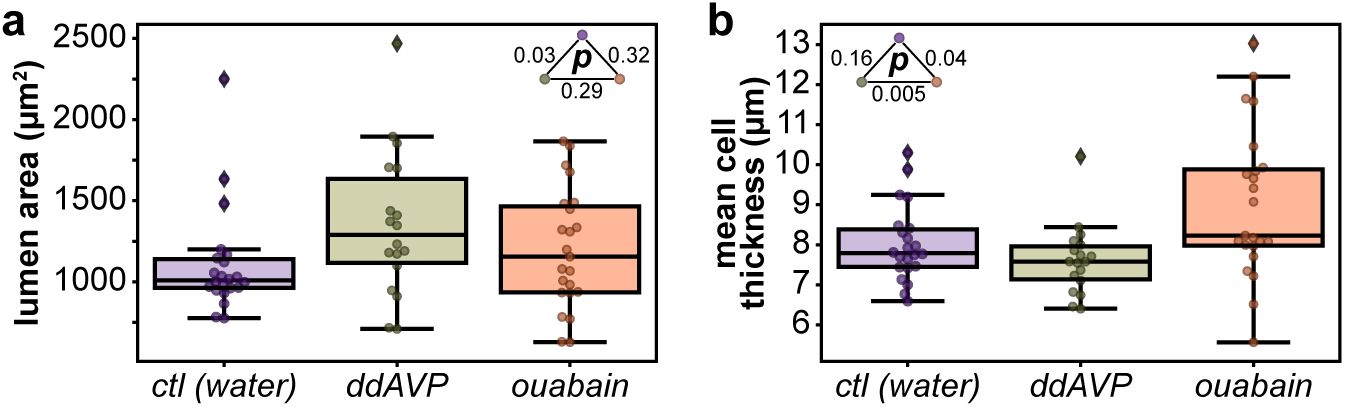
Quantification of MDCK spheroids grown in Matrigel for 7 days treated with ddAVP or ouabain for 4 hours. **a**, Quantification of cross-sectional luminal area in control, ddAVP, and ouabain conditions (p-values from rank-sum test). **b**, Quantification of mean cross-sectional cell wall thickness in control, ddAVP, and ouabain conditions (p-values from rank-sum test). Box plots in **a, b** show median, quartiles of dataset, and whiskers extending to maximum and minimum of distributions. For plots **a-b, *n*** = 22, 18, 23 spheroids for control, ddAVP, and ouabain conditions, respectively.

**Supplementary Figure 3.**
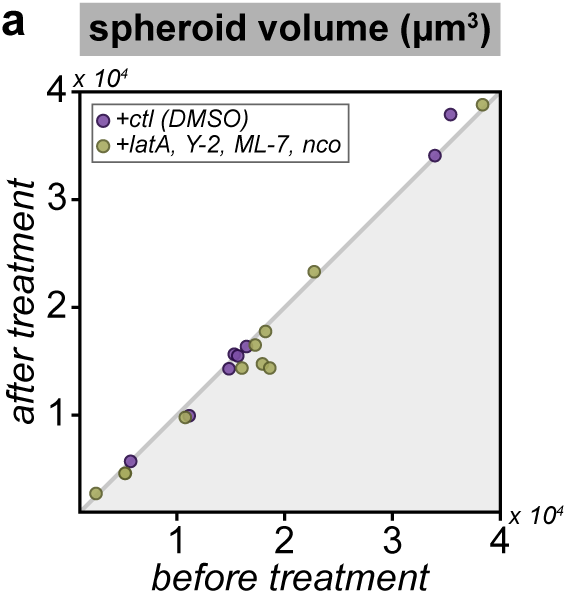
**a**, Plot of spheroid volume before (*t* = −2 min) and after (*t* = +18 min) treatment (rank-sum test DMSO-cytoskeletal inhibitors p = 0.13, Wilcoxon signed rank test *Δt*(DMSO) p = 1, *Δt*(cytoskeletal inhibitors) p = 0.02). ***n*** = 8 and 11 spheroids for DMSO and cytoskeletal inhibitors conditions, respectively.

**Supplementary Figure 4.**
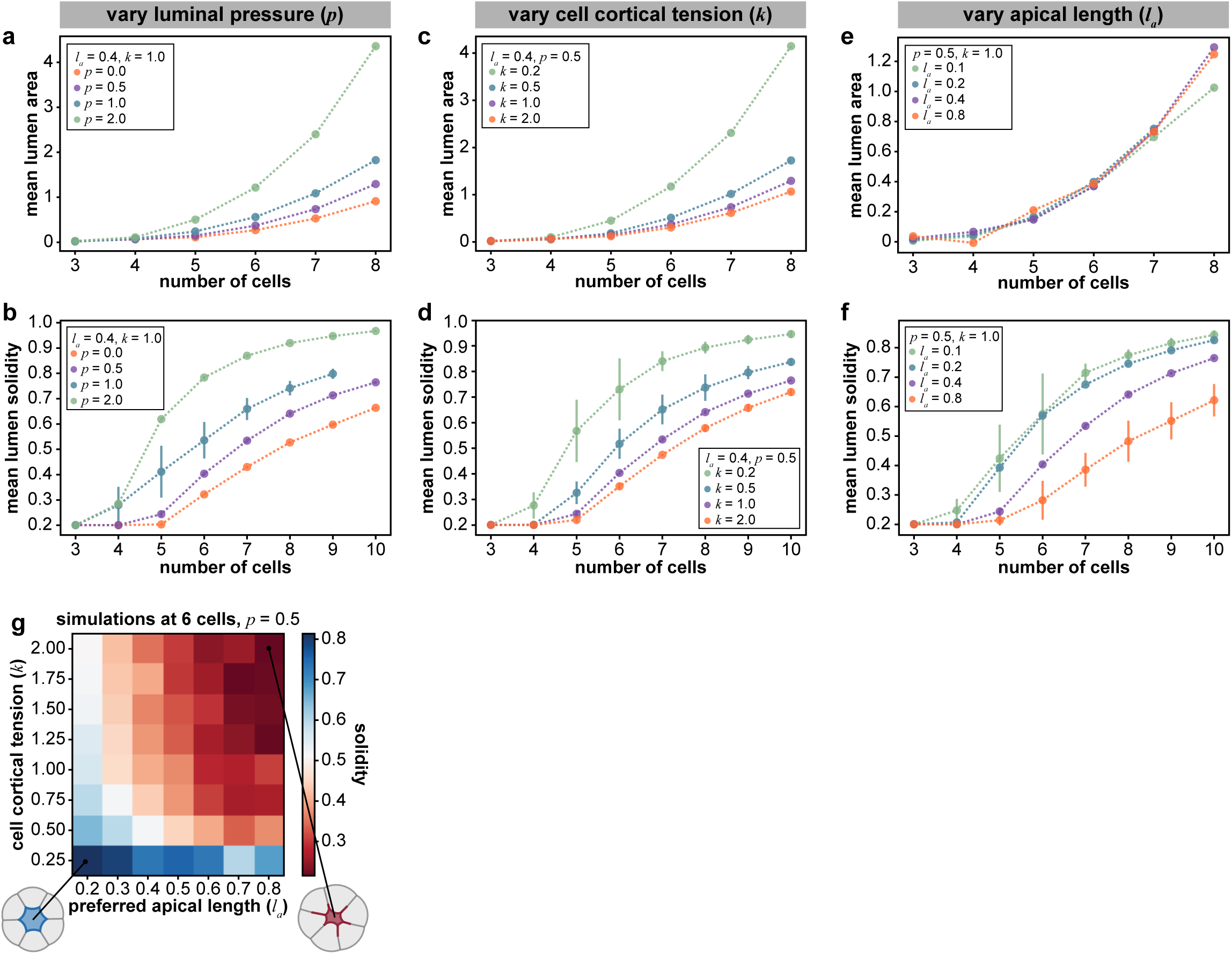
**a, c, e**, Plots of mean lumen area as a function of number of cells in a spheroid from simulations of 2D vertex-like model, keeping preferred apical length and cell cortical tension fixed (*l*_*a*_= 0.4, *k* = 1.0) while varying luminal pressure (*p* = 0.0-4.0) **(a)**, keeping preferred apical length and luminal pressure fixed (*l*_*a*_ = 0.4, *p* = 0.5) while varying cell cortical tension (*k* = 0.0-2.0) **(c)** and keeping luminal pressure and cell cortical tension fixed (*p* = 0.5, *k* =1 .0) while varying preferred apical length (*l*_*a*_ = 0.1-0.8) **(e). b, d, f**, Plots of mean lumen solidity as a function of number of cells in a spheroid from simulations of 2D vertex-like model, keeping preferred apical length and cell cortical tension fixed (*l*_*a*_ = 0.4, k =1 .0) while varying luminal pressure (*p* = 0.0-4.0) **(b)**, keeping preferred apical length and luminal pressure fixed (*l*_*a*_ = 0.4, *p* = 0.5) while varying cell cortical tension (*k* = 0.0-2.0) **(d)** and keeping luminal pressure and cell cortical tension fixed (*p* = 0.5, *k* =1 .0) while varying preferred apical length (*l*_*a*_ = 0.1-0.8) **(f). (g)** Heatmap of mean solidity for outputs of simulations while varying cell cortical tension and preferred apical length. Representative outputs of maximum (blue) and minimum (red) solidity reached in simulations.

**Supplementary Video 1** | Representative movies of MDCK spheroids expressing Lifeact-RFP treated with DMSO vehicle control (**left**) and cytoskeletal inhibitor cocktail (latrunculin A, Y-27632, ML-7, and nocodazole, **right**). Scale bar is 10 μm.

